# An *In Silico* Model of DNA Repair for Investigation of Mechanisms in Non-Homologous End Joining

**DOI:** 10.1101/318139

**Authors:** John W. Warmenhoven, Nicholas T. Henthorn, Marios Sotiropoulos, Nickolay Korabel, Sergei Fedotov, Ranald I. Mackay, Karen J. Kirkby, Michael J. Merchant

## Abstract

In human cells, non-homologous end joining is the preferred process to repair radiation induced DNA double strand breaks. The complex nature of such biological systems involves many individual actions that combine to produce an overall behaviour. As such, experimentally determining the mechanisms involved, their individual roles, and how they interact is challenging. An in silico approach to radiobiology is uniquely suited for detailed exploration of these complex interactions and the unknown effects of specific mechanisms on overall behaviour. We detail the construction of a mechanistic model by combination of several, experimentally supported, hypothesised mechanisms. Compatibility of these mechanisms was tested by fitting to results reported in the literature. To avoid over fitting, individual mechanisms within this pathway were sequentially fitted. We demonstrate that using this approach the model is capable of reproducing published protein kinetics and overall repair trends. This process highlighted specific biological mechanisms which are not clearly defined experimentally, and showed that the assumed motion of individual double strand break ends plays a crucial role in determining overall system behaviour.

## 1 Introduction

Radiotherapy is a well-established treatment modality for cancers, with substantial clinical experience in delivering palliative and radical photon based procedures. Proton beam therapy is a fast developing alternative modality which offers advantages for specific cancer sites due to the ions stopping a defined depth into the patient. Protons also produce a different biological effect for the same dose delivered, parameterised as the proton Relative Biological Effectiveness (RBE); defined as the ratio between proton and photon doses that are required to cause an equivalent amount of cell killing. Current clinical practice is to scale proton dose by a fixed RBE of 1.1 (DeLuca *et al*, 2007).

However, experimental data shows a large variance in RBE depending on parameters describing both the tissue environment and the beam quality (Paganetti *et al*, 2002; Paganetti, 2014; IAEA & ICRU, 2008). Studies show that a variable RBE can increases not only the biologically effective range, but also the biologically effective dose at the distal edge of the proton beam (Marshall *et al*, 2016). Consequently, studies based on *in vitro* data have shown the possibility of these biological uncertainties to negatively impact clinical proton treatment plans (Wedenberg & Toma-Dasu, 2014; McNamara *et al*, 2016; Tilly *et al*, 2005; Carabe *et al*, 2012; Wedenberg *et al*, 2013). Tilly *et al.* and McNamara *et al.* demonstrate the possibility of increased normal tissue complication probability (NTCP) whilst Wedenberg *et al.* demonstrate the possibility of a lower tumour control probability (TCP). Carabe *et al.* show that there are clinical situations in which the relatively small biological range uncertainty is relevant. Development of future treatment planning systems should therefore include RBE optimisation to further exploit the inherent physical advantages of proton beam therapy (Giovannini *et al*, 2016; Cao *et al*, 2015). Computational models are an efficient way of probing various biological and physical mechanisms in order to illuminate their possible relationships to RBE.

Several *in silico* models of DNA damage and repair have been proposed that assume a random spread of breaks, homogeneous throughout the cell or centred around a track (Sachs *et al*, 2000; Ballarini & Ottolenghi, 2004; Eidelman *et al*, 2012). An alternative methodology is to analyse radiation track structure interactions within the cell nucleus (Bernal *et al*, 2015). This provides the specific geometrical distributions of double strand breaks within the nucleus caused by radiations of different types and qualities, therefore making it possible to investigate how changes in proximity of breaks affects the repair machinery (Moore *et al*, 2014). Furthermore, it also enables explicit inclusion of repair retarding complexities such as base lesions and additional single strand breaks (Schipler & Iliakis, 2013). Due to the small number of double stand breaks at low doses of ionising radiation, discrete stochastic models are most suited to simulating the repair of these damages (Friedland & Kundrát, 2013; Friedland *et al*, 2010b). Such models also allow for inclusion of specific mechanistic interpretations of the interactions between the repair machinery and the complexity or proximity of double strand breaks; something that is difficult with more top down descriptions. Therefore combining discrete, stochastic, mechanistic models with damage derived from detailed track structure simulations would produce a model suitable for studying the broadest range of radiation doses. Thorough investigation of the emergent behaviour of such a system could give insight into the operation of cellular repair machinery and subsequently into the experimentally observed variations in RBE.

DNA double strand breaks have multiple pathways of repair, the most dominant being Non-Homologous End Joining (NHEJ) (Chiruvella *et al*, 2013) and Homologous Recombination (HR) (Jasin & Rothstein, 2013). The non-homologous end joining pathway is available throughout the cell cycle, and is the only pathway available outside of the replication phases (Thompson, 2012; Brown *et al*, 2017). Furthermore, the most severely radio-sensitive phenotypes have mutations of non-homologous end joining genes (Thompson, 2012) showing that this pathway plays a dominating role in cell survival. Initial focus on this pathway is therefore justifiable and facilitates easy designing of *in vitro* verification experiments that isolates the action of non-homologous end joining. However, the complexity of biological systems, even when reduced to a single pathway, presents a significant challenge to *in vitro/vivo* experiments when attempting to unambiguously define mechanisms responsible for DNA repair. This has resulted in separate publication of many suggested mechanisms within the NHEJ pathway that are not contradicted by experimental evidence. These mechanisms, although individually plausible, are not necessarily capable of reproducing observed behaviours of the entire cellular system when combined.

In this work we construct a composite system to model non-homologous end joining through the combination of several such published mechanisms. This mechanistic, Monte Carlo based model is developed within the Geant4-DNA tool-kit (Incerti *et al*, 2010; Bernal *et al*, 2015; Karamitros *et al*, 2014) which allows for event-by-event tracking of individual double strand break ends. Initial conditions were set from interfacing with a nano-dosimetric DNA damage model, directly linking radiation track structure to biological effect (Henthorn *et al*, 2017, 2018). The model produces as its output predictions of biologically measurable end points, including repair protein recruitment kinetics and double strand break repair kinetics, which are compared to published experimental data to assess the feasibility of the combined mechanisms.

## 2 Materials and Methods

### 2.1 Construction of Initial Model Conditions

The repair model takes as its input data in a standard DNA damage phase space format developed in collaboration with Queen’s University Belfast and Harvard Medical School. Damage models can utilise this standard to populate a phase space that can be passed either in the form of an ASCII file or directly if the damage model is integrated into the Geant4 simulation. In this work we use simulation results from the model described by Henthorn *et al.* (Henthorn *et al*, 2017, 2018) as input. This nano-dosimetric track structure model provides the geometric locations and detailed structure of DNA double strand breaks after irradiation of a nucleus by a specified beam. Individual double strand breaks are processed and separated into two exposed DNA ends at the closest pair of backbone breaks on opposite strands. Remaining backbone and base damages are assigned to individual break ends based on their positions relative to the separation site.

### 2.2 Implementation of Double Strand Break Objects

We have implemented double strand breaks using the existing Geant4-DNA mechanism for handling chemistry objects; that is an object consisting of a track and associated molecule. This allows use of the reaction and diffusion functionality of the Geant4-DNA Chemistry module (Karamitros *et al*, 2014). To monitor the state of each individual double strand break object as the system evolves a new software class is implemented and associated to the track as auxiliary track information. This class is a data structure storing the number of single strand breaks and base lesions (see section 2.1), break ID and correct partner ID (see section 2.4), waiting and diffusion times (see section 2.3), and starting location (see section 2.5) associated with each break end object. There are also empty copies of these variables for use when two break end objects react to form one synaptic object (see section 2.4.3).

### 2.3 Motion

It has been experimentally demonstrated that double strand breaks move by sub-diffusion (Girst *et al*, 2013; Lucas *et al*, 2014; Miné-Hattab *et al*, 2017), and, although not explicitly concluded, there is experimental evidence that this applies to individual double strand break ends (Soutoglou *et al*, 2007). To the best of our knowledge this type of motion has not yet been explicitly included in any mechanistic *in silico* models of DNA repair. There are different microscopic mechanisms that lead to the emergence of sub-diffusive motion (Sokolov, 2012). In this work we have chosen to implement a continuous time random walk (CTRW) model of sub-diffusion (for a detailed discussion of anomalous diffusion in general see Metzler *et al.* (Metzler & Klafter, 2000)). This represents the double strand break ends undergoing transient states of confined motion, such as would be expected in a system with rapid formation and dissociation of synaptic complexes (Graham *et al*, 2016). To implement such a continuous time random walk model double strand break objects are assigned a waiting time drawn from:

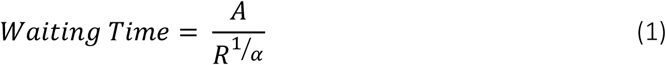

Where R is a uniform random number between 0 and 1, A is the minimum waiting time and α is the anomalous diffusion exponent. In this work the anomalous diffusion exponent was set to 0.5 and the minimum waiting time was set to 1 ms in order to generate the waiting times. Whilst a double strand break object has a waiting time associated it is trapped and cannot diffuse. Waiting times are tracked using the internal clock of the simulation and once reduced to 0 seconds the double strand break object is released. Diffusion then occurs through normal Brownian motion means of stochastic displacement by a distance drawn from a normal distribution. The distribution is described by the diffusion coefficient and the time the object is allowed to diffuse. In our model the diffusion time is set at 1 picosecond, which promotes a single ‘jump’ event without interference of any chemical processes. The ‘jump diffusion coefficient’ is taken to be the free fitting parameter to manipulate the overall scale of motion and fit to literature data. In this way we have implemented a mechanism which can lead to sub-diffusive motion. The mechanism is linked to transient confinement of double strand break ends such as could be explained by alternating association/dissociation processes (Saxton, 2007).

### 2.4 Time Evolution

#### 2.4.1 Implementation of the Non-Homologous End Joining Process

Time evolution of individual double strand break ends is governed by a series of time constant based state changes (see section 2.4.2). The states are linked according to the scheme outlined in Figure 1. This represents the progressions through the canonical non-homologous end joining pathway.

**Figure 1:**
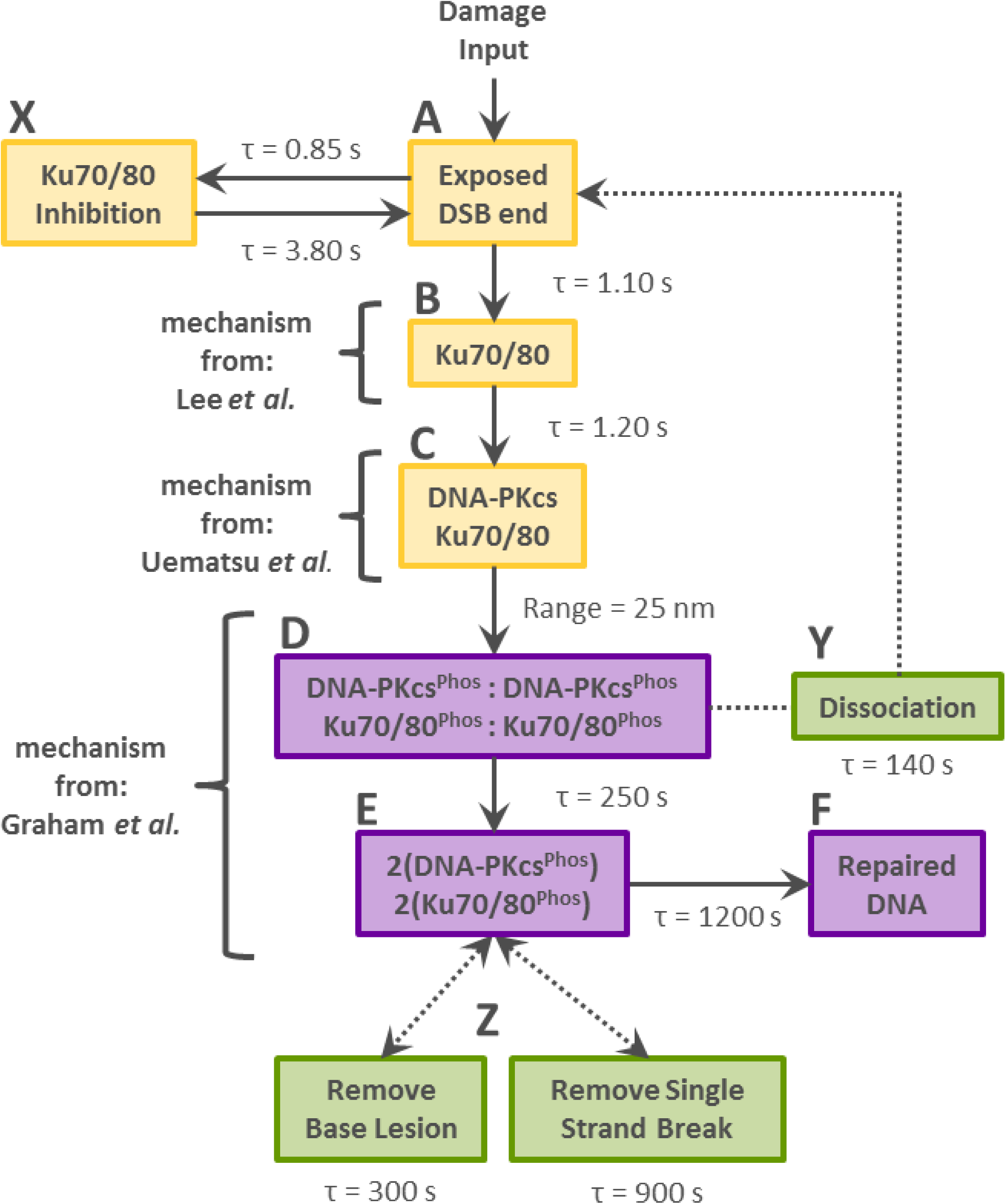
Scheme of the non-homologous end joining pathway implemented in this work with associated time constants. Boxes A-F and X represent the different states double strand break objects can occupy in the simulation, boxes Z and Y and arrows represent the different mechanisms which can change the state of a double strand break object in the simulation. States A-C and X represent a single double strand break end; states D-F represent a synaptic complex of two double strand break ends. Progress between states A-C, A to X, D-F, and D to A is governed by a time constant based stochastic process. Progress from states C to D is governed by a proximity based reaction which produces a single synaptic complex from two double strand break ends. Process Y splits this synaptic complex back into two individual double strand break ends. Also noted in the figure are the areas of influence of the three experimentally supported mechanisms combined by this work. Lee *et al.* suggested that Ku70/80 does not dissociate until phosphorylated, Uematsu *et al.* suggesting DNA-PKcs does not dissociate until phosphorylated, and Graham *et al.* showing formation of transient synaptic complexes between DNA-PK complexes.

State A represents an exposed double strand break end which may have associated single strand breaks and base lesions as discussed in section 2.2. Recruitment of the Ku70/80 heterodimer is widely reported in literature to be the first step in the canonical non-homologous end joining pathway (Ma *et al*, 2004; Lieber, 2010; Chiruvella *et al*, 2013; Wang & Lees-Miller, 2013; Brown *et al*, 2017) (State B). Ku70/80 cannot dissociate from DNA ends until phosphorylated (mechanism proposed by Lee *et al.* (Lee *et al*, 2015)). Double strand break ends may alternatively become temporarily inhibited towards recruitment of Ku70/80 (state X). The inhibition is a catch all state which represents competition from other proteins with affinities for exposed double strand break ends. Following attachment to a double strand break end, Ku70/80 recruits the DNA-Protein Kinase catalytic subunit (DNA-PKcs) to form the DNA-PK complex (state C). DNA-PKcs cannot dissociate from DNA ends until phosphorylated (mechanism proposed by Uematsu *et al.* (Uematsu *et al*, 2007)). DNA-PK is required for the formation of synaptic complexes between proximal double strand break ends (Graham *et al*, 2016) (state D, see section 2.4.3).

Once in a synaptic complex DNA-PKcs autophosphorylates, inducing conformational changes necessary for further processing and allowing it to phosphorylate Ku70/80 (Uematsu *et al*, 2007; Lee *et al*, 2015). This initial “long range” synaptic complex is prone to dissociation (mechanism proposed by Graham *et al.* (Graham *et al*, 2016)) and due to the phosphorylated state of both Ku and DNA-PKcs the entire complex dissociates to state A (Uematsu *et al*, 2007; Lee *et al*, 2015) (process Y, see section 2.4.3). If the complex survives, progression to a stable “short range” synaptic complex follows (state E) (mechanism proposed by Graham *et al.* (Graham *et al*, 2016)). Processing of the break site occurs in state E, repairing all base lesions and extra DNA backbone lesions present (processes Z, see section 2.4.2) before allowing final ligation to state F.

#### 2.4.2 State Change

State changes are implemented as custom Geant4 processes that change the molecule definition associated with the track. Seven different definitions are created to represent the different states a double strand break object can occupy. These are states A-F and X from Figure 1. The time constants used in these processes represent both the attachment and action of specific repair proteins. Cleaning of complexities, state Z in Figure 1, are special cases of the above processes. On execution no change is made to the object state. Instead the number of base lesions or single strand breaks associated with the synaptic complex is reduced. The specific double strand break end this applies to within the complex is determined randomly. Transition from state E to F cannot successfully complete unless all associated base lesions and single strand breaks have been removed.

#### 2.4.3 Reactions and Failure after Synapse

Both DNA-PK mediated and XRCC4 mediated synaptic complex formation for complex and simple double strand breaks respectively have been reported (Reid *et al*, 2015; Graham *et al*, 2016; Li *et al*, 2014). In this work we have chosen to explicitly model only the DNA-PK mediated synaptic complex formation as suggested by Graham *et al.* (Graham *et al*, 2016) due to available experimental data to fit to. Synaptic complex formation is treated as a chemical reaction utilising the provided functionality of Geant4-DNA. Two DNA-PK objects within 25 nm (Friedland *et al*, 2010a) of each other react to form a single synaptic complex object. Outside of this range the reaction probability is governed by the Geant4-DNA diffusion controlled Brownian bridge process (Karamitros *et al*, 2014) modified to accommodate sub-diffusive motion. A reaction replaces two double strand break end objects with a single synaptic complex object. The state information of both double strand break end objects is stored together (see section 2.2) and associated with the new synaptic complex object. This preserves the information required for final data gathering (see section 2.5) and dissociation of the complex. Dissociation replaces the single synaptic complex object with two double strand break end objects and merging of the associated state information is reversed.

### 2.5 Data Gathering

Recruitment kinetics of involved repair proteins and overall double strand break repair kinetics are extracted by real time tracking of object states. Simulations are repeated in order for the stochastic state changes to converge on an average result. Residuals double strand breaks are scored as the initial number of double strand breaks reduced by the number of fully ligated ‘fixed’ breaks in the system at that time point. This represents all double strand break ends which have become isolated or are still undergoing repair. Data is normalised for each individual simulation and then combined over all repeats to give the average behaviour for a given beam quality. Displacement of all double strand break objects from their initial location is tracked in one second intervals for the first 300 seconds to calculate the mean squared displacement. Misrepair events are scored at the end of each individual simulation by comparing identities of double strand break ends in synaptic complexes.

## 3 Results

### 3.1 Recruitment Kinetics of DNA-PKcs and Ku70/80

The stoichiometry of Ku70/80 and DNA-PKcs is one each per double strand break end. A high density of breaks is therefore required for these proteins to form visible foci under most microscopy conditions. To meet this requirement Uematsu *et al.* (Uematsu *et al*, 2007) reported on 365 nm pulsed nitrogen laser generated double strand breaks, stating that they created 2500-3700 per 1.7 μm^2^ spot. These breaks are also considered to be complex in nature and thus require the DNA-PKcs mediated NHEJ pathway described in this work (Li *et al*, 2014). Uematsu *et al.* also report experimental data from uranium ions accelerated to 3.8 MeV/u. The high LET (>12,800 keV/μm) of these ions is expected to result in similarly highly clustered and complex damage.

Figure 2A and 2B shows good agreement between the simulated behaviour and literature reported experimental data from cell lines derived from Chinese hamster ovary (CHO) cell lines, namely Xrs6 and V-3, regardless of radiation used. Initial rapid recruitment of Ku70/80 and DNA-PKcs reaches 50% of maximum in 2-3 seconds before slowing, reaching 80% at ^~^10 seconds. Data from non-CHO cell lines show slightly more rapid recruitment of Ku70/80 which is not reproduced as well. Andrin *et al.* (Andrin *et al*, 2012) use human bone osteosarcoma epithelial cells (U2OS) in their work and Hartlerode *et al.* (Hartlerode *et al*, 2015) show similar Ku70/80 recruitment in mouse embryonic fibroblast stem cells (MEF). Fitting the DNA-PKcs recruitment to experimental data was trivial; requiring manipulation of only the state B to state C time constant (Figure 1) to produce a slight delay on the Ku70/80 recruitment resulting in the correct behaviour. Obtaining the correct behaviour of Ku70/80 recruitment was more involved, requiring manipulation of the 3 relevant time constants, with inhibition (A to X, Figure 1) playing an important role. Final time constants selected are Ku-inhibition = 0.85 seconds, release from inhibition = 3.8 seconds, Ku70/80 recruitment = 1.1 seconds, DNA-PKcs recruitment = 1.2 seconds.

**Figure 2:**
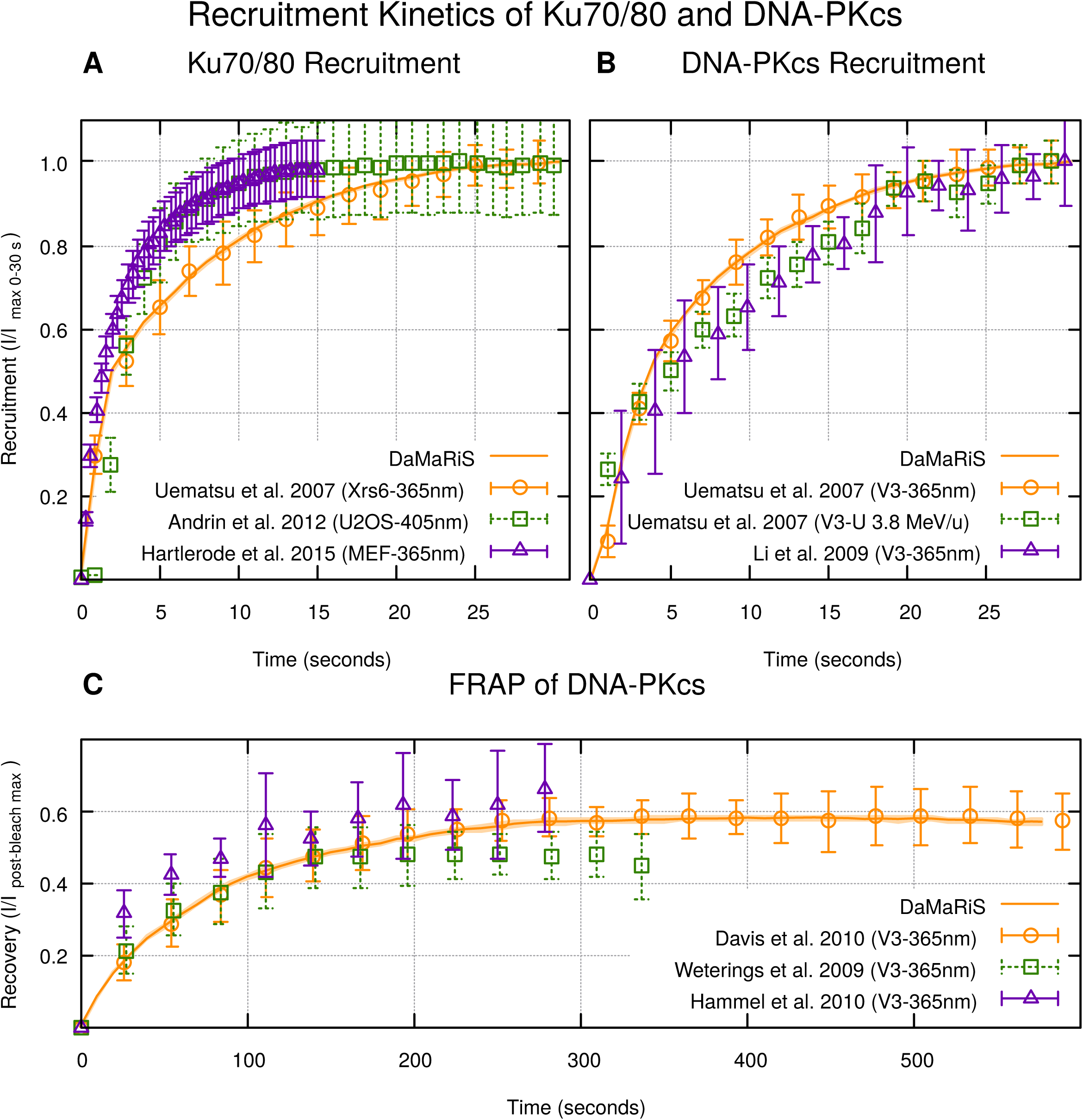
Recruitment of (A) Ku70/80 and (B) DNA-PKcs to exposed double strand break ends; values are normalised to the maximum value achieved during the 30 seconds plotted. Individual simulation repeats were normalised to their initial number of double strand breaks. Final values are the average of these normalised trends. (C) Recovery of fluorescent DNA-PKcs at double strand break ends after a simulated photo-bleaching event; intensities are set to 0 at the photobleaching event and normalised to the maximum value achieved during the 600 seconds plotted. To remove the influence of a variable number of initial double strand breaks, individual simulation results were normalised to themselves before all repeats were averaged to produce final values. The line is the simulation results and empty symbols are the corresponding experimental results. Shaded area around the line is the standard error in the mean of 200 repeats. Empty symbols are experimental values taken from the referenced papers with brackets indicating the cell line followed by the radiation type used. Error-bars associated with Hartlerode *et al.*(Hartlerode *et al*, 2015) data are standard errors in the mean, the source of error-bars associated with Weterings *et al.*(Weterings *et al*, 2009) data is unknown, and all other error-bars are the reported standard deviation.

**Table 1:** Summary of the reduced χ^2^ for the different experimental datasets compared to the simulated behaviour. Data from the different datasets had different measuring time-points and consequently exactly corresponding simulation time points were not available. Experimental data was therefore compared to the nearest simulation point within a 1% time deviation where available. Two values are given for Hartlerode *et al.* data as the method the authors used to calculate SEM was not clear (see supplementary material in Hartlerode *et al.* (Hartlerode *et al*, 2015)). We back calculated the standard deviation either assuming the 88 cells were used to calculate SEM, or instead that the three experiments were used to calculate SEM, given in brackets.

| Authors | Protein | Reduced $\chi^2$ | Max / Average / $\sigma$ of Time Deviation (%) |
| --- | --- | --- | --- |
| Uematsu *et al.* | Ku70/80 | 0.020 | 0.74 / 0.17 / 0.19 |
| Andrin *et al.* | Ku70/80 | 0.562 | 0.96 / 0.15 / 0.21 |
| Hartlerode *et al.* | Ku70/80 | 0.069 (1.336) | 0.72 / 0.19 / 0.21 |
| Uematsu *et al.* (LASER) | DNA-PKcs | 0.033 | 0.95 / 0.51 / 0.28 |
| Uematsu *et al.* (Uranium) | DNA-PKcs | 1.167 | 0.82 / 0.48 / 0.17 |
| Li *et al.* | DNA-PKcs | 0.577 | 0.92 / 0.34 / 0.31 |

The simulated fluorescent recovery after photobleaching (FRAP) behaviour in Figure 2C is generated by a customisation of the simulation pathway to include photobleached counterparts of components that involve molecules of DNA-PKcs. At 30 seconds a bleaching event is simulated converting all double strand break ends with attached DNA-PKcs into photobleached versions. Recovery is measured as the rate at which these bleached ends dissociate and acquire new unbleached DNA-PKcs. The behaviour is again compared to literature reported laser generated double strand break data for the same reasons as described above. Weterings *et al.* (Weterings *et al*, 2009) agrees with the theoretical fit to simulated data up until 140 seconds (Inline). After this the fit worsens (Inline)as the data plateaus earlier in spite of the same cell line being used. The lack of data from Hammel *et al.* (Hammel *et al*, 2010) for times greater than 300 seconds prohibit identification of a plateau.

The fluorescent recovery of a system can be diffusion limited, reaction limited, or a complex mix of the two. Diffusion controlled systems are limited by the rate at which non-bleached proteins diffuse into the bleached area and replace bleached proteins. The behaviour of these systems tends towards the equation (Soumpasis, 1983):

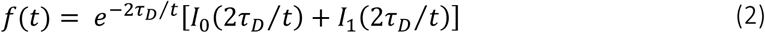

where f(t) is the relative fluorescence, t is the time, and τ_D_ is the “characteristic” diffusion time. Reaction controlled systems are limited by the rate at which bleached proteins detach from binding sites in the breached region, allowing non-bleached proteins to replace them. The behaviour of these systems tends towards the equation (Bulinski *et al*, 2001):

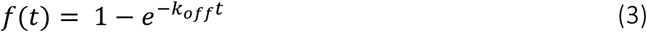

where f(t) is the relative fluorescence, t is the time, and k_off_ is the detachment rate of the protein in question. Fitting these equations to the Davis *et al.* data, Equation 2 with τ_D_ = 33.46 seconds gives a reduced χ^2^ of 0.595 whilst Equation 3 with k_off_ = 1.26 ×10^−2^ s^−1^ gives a reduced χ^2^ of 0.025. From this it can be seen that the Davis *et al.* data is much better fitted by a reaction limited, rather than diffusion limited system. Due to the rapid recruitment of Ku70/80 and DNA-PKcs derived from the data in Figure 2A and 2B, the dissociation time constant is the remaining parameter which has a dominating influence on the initial behaviour of the recovery curve for the first 300 seconds in our model. The final time constant selected for dissociation of transient synaptic complexes is 140 seconds.

**Table 2:** Summary of the reduced χ^2^ for the different experimental datasets compared to theoretical fits assuming diffusion limited or reaction limited recovery. All works are fitted best by assuming reaction limited recovery apart from that of Hammel *et al.*.

|  | Diffusion Limited |  | Reaction Limited |  |
| --- | --- | --- | --- | --- |
| Authors | $\tau_D (s)$ | $\chi^2_v$ | $k_{off} (s^{-1})$ | $\chi^2_v$ |
| This work | 37.78 | 0.263 | 0.0128 | 0.008 |
| Davis *et al.* | 34.37 | 0.595 | 0.0125 | 0.025 |
| Weterings *et al.* | 20.81 | 0.281 | 0.0206 | 0.022 |
| Hammel *et al.* | 32.00 | 0.070 | 0.0169 | 0.379 |

In the simulation, DNA-PKcs is assumed to be retained until the double strand breaks are converted into a fully fixed state. Due to the complex nature of all the breaks, this state is only reached after backbone breaks and base lesions are cleaned. The plateau of the recovery curve from 300-600 seconds in Figure 2C therefore gives a lower bound for the time constants associated with these actions. Final time constants selected are Backbone Clean-up = 300 seconds, Base Lesion Clean-up = 900 seconds in agreement with those of Friedland *et al.* (Friedland *et al*, 2010a).

The fluorescent recovery plateaus at ^~^60% of pre-bleaching event intensity as observed by Davis *et al.*. In the simulation DNA-PKcs can only dissociate as part of the entire synaptic complex dissociating. Dissociation of these synaptic complexes can only happen whilst they are in their unstable, transient, states. Therefore, manipulation of the rate at which synaptic complexes stabilise is sufficient to vary the amount of bleached DNA-PKcs trapped at double strand breaks and determine the overall recovery. The final time constant selected for stabilisation of transient synaptic complexes is 250 seconds.

### 3.2 Repair Kinetics

DNA damage patterns were generated using the model published by Henthorn *et al.* (Henthorn *et al*, 2017, 2018) with beam parameters corresponding to those used experimentally by Chaudhary *et al.* (Chaudhary *et al*, 2016). This allowed comparison of high (13.7 keV/μm) and low LET (1.77 keV/μm) proton beams directly to experimental data (Figure 3) in order to fit the remaining final ligation time constant and the parameters that govern the mobility of double strand break ends. The primary effect of the final ligation time constant is to determine the delay in the sharp drop off in residuals observed in the first 100 minutes in Figure 3 but has minimal impact on the slope of this curve. The initial experimental time points at 30 minutes therefore set an upper limit to the final ligation time constant in order to avoid overshooting them. The final time constant selected for final ligation is 1200 seconds.

**Figure 3:**
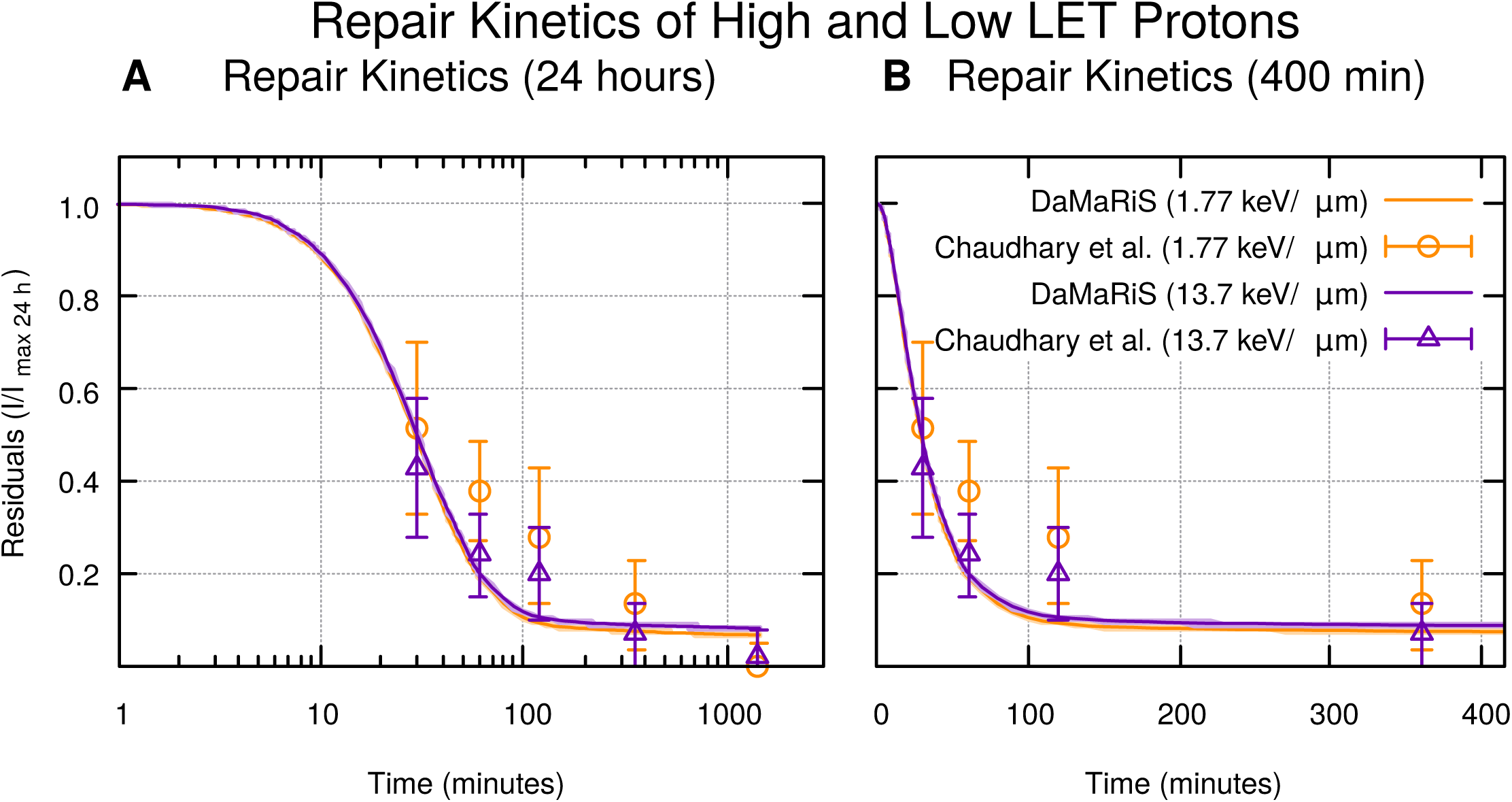
Resolution of double strand breaks over (A) a 24 hour period and (B) a zoom in of the first 400 minutes, for two separate LET proton radiations. Residuals were calculated as the initial number of double strand breaks reduced by the number of fully ligated ‘fixed’ breaks in the system at that time point. Individual simulation repeats were normalised to their initial number of double strand breaks. Final values are the average of these normalised trends. Lines are simulation results with the shaded area around the line representing the standard error in the mean of 200 repeats. Empty circles are the corresponding experimental results of 53BP1 foci resolution from Chaudhary *et. al*(Chaudhary *et al*, 2016), with error-bars representing the reported standard deviations.

The mechanisms of the repair machinery have thus far been determined by fitting to experimental data which has resulted in limiting the period of action to the initial two hours. Consequently, to fit our model to the residual breaks at 24 hours we must address the diffusion of double strand break ends.

### 3.3 Diffusion

The yield of residual double strand breaks at 24 hours was scored for 1 Gy of 13.72 keV/μm proton irradiation whilst varying the magnitude of the jump diffusion coefficient assigned to double strand break ends. The average number of residual breaks reported by the simulation was compared with the experimental data of the corresponding time point. The best fit was achieved (^~^4.6 residual foci (Chaudhary *et al*, 2016)) at a jump diffusion coefficient of 2.8×10^11^ nm^2^/s. The residual double strand break yield produced by this motion was then compared across a range of LET to residual 53BP1 foci experimentally observed by Chaudhary *et al.* (Chaudhary *et al*, 2016), Figure 4A. Our model reproduces a similar trend in LET dependence with gradients for both fitted lines within error. There is however a small systematic over-estimation shown by the difference in intercepts of the fitted lines of 0.56±0.35 breaks.

**Figure 4:**
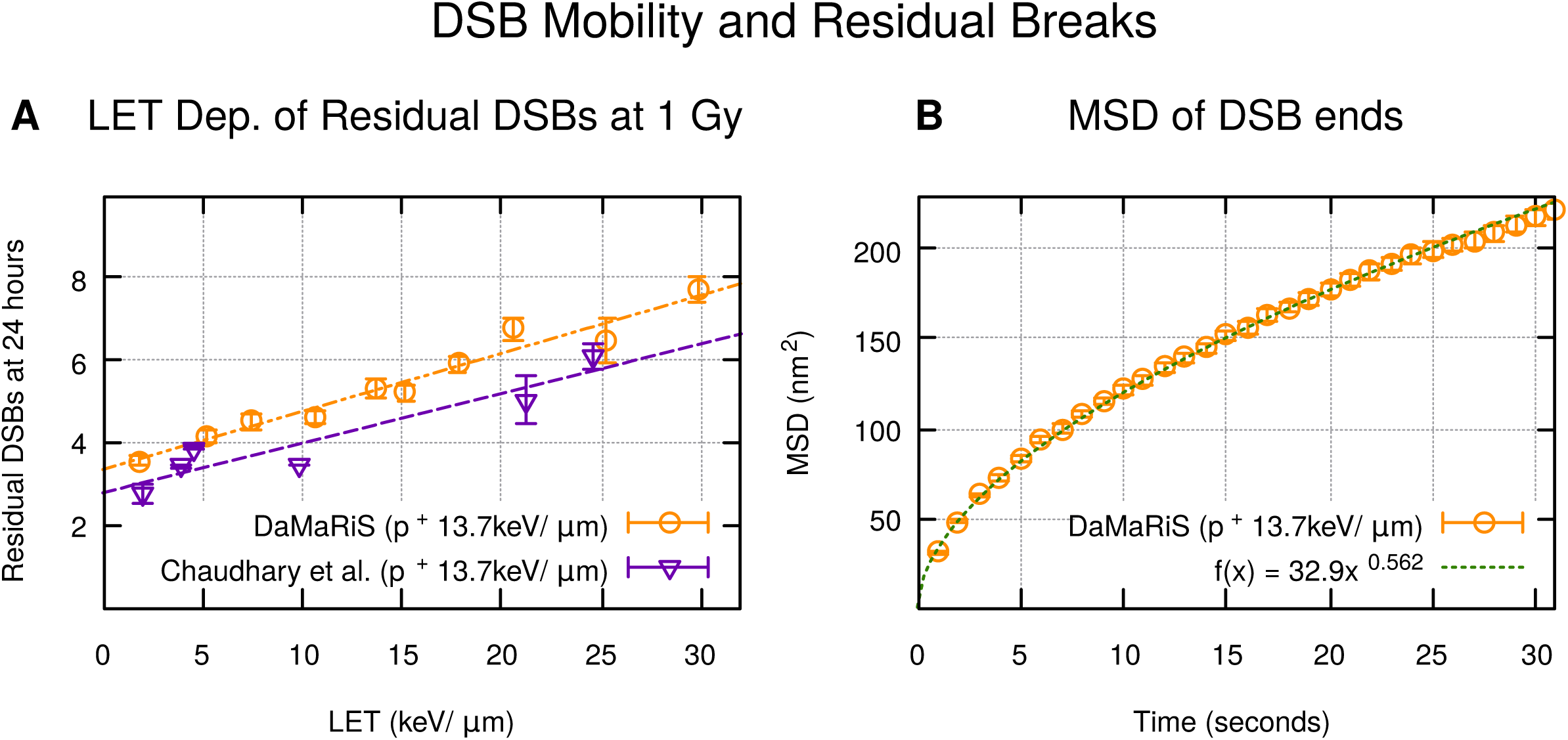
(A) Residual breaks at 24 hours for 1 Gy of proton radiation at a range of LET showing the same trend as experimental measurements. Empty circles are simulation results, with error-bars showing the standard error in the mean of 69 repeats for the ^~^25.3 keV/μm point and 200 for the rest. Experimental values are shown as empty triangles and taken from Chaudhary *et. al.*(Chaudhary *et al*, 2016), with error-bars representing the standard error in the mean. Dashed lines are linear fits. Fit to this work has m=0.141±0.011 and c=3.34±0.18, fit to Chaudhary *et al.* data has m=0.121±0.021 and c=2.78±0.30. (B) Mean squared displacement of double strand breaks caused by 1 Gy of 13.72 keV/μm protons over the initial 30 seconds. Empty circles are simulation results with error-bars representing the standard error in the mean of 200 repeats. The dashed line is Equation 4 fitted with values of D_α_ = (32.9±0.4) nm^2^/s^0.56^, and α = 0.562±0.004.

To quantify the sub-diffusive behaviour in the model, the mean squared displacement of double strand break ends were tracked over the initial 300 seconds of the simulation for 200 repeats, Figure 4B. The data was fitted with a power law:

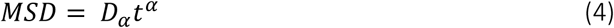

with D_α_ in units of nm^2^/s^α^ being the general diffusion coefficient, and α being the anomalous diffusion exponent. α = 1 indicates normal Brownian diffusion following Fick’s law, α > 1 indicates super-diffusive properties, and α < 1 indicates sub-diffusion.

Mobility of individual double strand break ends are difficult to investigate, however, there are numerous groups that have investigated the mobility of double strand break loci or chromosomal loci. Cabal *et al.* (Cabal *et al*, 2006) measured the motion of fluorescently labelled GAL genes in *Saccharomyces cerevisiae*, measuring α = 0.42±0.01 for gene foci not confined to the nuclear envelope. Weber *et al.* (Weber *et al*, 2010) studied fluorescently labelled chromosomal loci in *E. coli* and *Vibrio cholerae*, measuring α to range from 0.35±0.02 to 0.44±0.06 for various gene foci. Bronstein *et al.* (Bronstein *et al*, 2009) used GFP-tagged TRF2 in U2OS cells to measure α for telomeres, observing growth from 0.32±0.12 to 0.51±0.20 during initial anomalous motion. Girst *et al.* (Girst *et al*, 2013) used GFP-tagged MDC1 in U2OS cells to measure0020α = 0.49±0.05 for double strand break foci after carbon irradiation and α = 0.50±0.05 after proton irradiation. Girst *et al.* did not observe splitting of foci and therefore concluded that individual double strand break ends must have a similar scaling of MSD with time. The measured value of the anomalous diffusion exponent, α, for individual double strand break ends in this work is 0.562±0.004 (Figure 4B). This agrees well with the work of Girst *et al.*.

In contrast to the relatively similar measurements of α, measurements of the general diffusion coefficient by the same groups differ considerably. Cabal *et al.* reported coefficients on the order of ×10^−2^ μm^2^/t^α^ for chromosomal foci in yeast, Bronstein *et al.* reported coefficients on the order of ×10^−3^ μm^2^/t^α^ for telomeres in bacteria, and Girst *et al.* reported coefficients on the order of ×10^−4^ μm^2^/t^α^ for double strand break foci in human cells. The measured value of the general diffusion coefficient, D_α_, for individual double strand break ends in this work is (32.9±0.4) nm^2^/s^0.56^, on the order of ×10^−5^ μm^2^/t^α^. This is an order of magnitude smaller than the motion measured by Girst *et al.*, indicating tighter confinement of the constituent ends than the parent double strand break.

## 4 Discussion

The complex nature of biological systems involves many individual actions that combine to produce an overall behaviour. To model these systems *in silico*, either a top down or bottom up approach can be used. Top down approaches, such as phenomenological models, have the advantage of limiting the number of variables used to recreate the observed behaviour of a system. Such an approach is useful for parameterising dependencies of the system on known variables, but does not provide clear insight into the mechanisms responsible. Bottom up approaches have the advantage of being useful in exploring the unknown effects of specific mechanisms on overall behaviour. As such they are useful tools for determining the compatibility of separately proposed, experimentally supported mechanisms. In this work we have combined several such mechanisms into one system and tested the subsequent overall behaviour against experimentally observed trends. We show that, with reasonable parameters for individual mechanistic steps, experimental results can be reproduced for relevant time points when combining mechanisms proposed by Graham *et al.* (Graham *et al*, 2016), Lee *et al.* (Lee *et al*, 2015), and Uematsu *et al.* (Uematsu *et al*, 2007). However, we show that a CTRW interpretation of the sub-diffusive motion of double strand break ends, as suggested by Bronstein *et al.* (Bronstein *et al*, 2009) and Girst *et al.* (Girst *et al*, 2013), is likely insufficient to explain overall repair kinetics.

Mechanistic, bottom up models must assign parameters to each mechanism which increases the danger of over-fitting. To guard against over-fitting, we have chosen to fit the model detailed in this work progressively, starting from fitting the earliest mechanisms in isolation and building from there. Table 3 summarises these results and other parameters used in the model.

**Table 3:** Table summarising the parameters used in this model. Values preceded by ‘Set at’ are not fitted parameters. The table is divided such that parameters manipulated concurrently to fit literature data are within the same cell. From this it can be seen that at most 3 parameters were used to reproduce any one experimentally observed behaviour.

| Process | Value | Fitted |
| --- | --- | --- |
| Ku70/80 recruitment | 1.1 s | Figure 2A |
| Ku70/80 inhibition | 0.85 s |  |
| Release from Ku70/80 inhibition | 3.8 s |  |
| DNA-PKcs recruitment | 1.2 s | Figure 2B |
| Reaction range to form synaptic complex | Set at 25 nm | Section 2.4.3 |
| Dissociation of synaptic complex | 140 s | Figure 2C |
| Clean Base Lesion | 300 s | Figure 2C |
| Clean Backbone Break | 900 s |  |
| Stabilisation of synaptic complex | 250 s | Figure 2C |
| Final ligation steps | 1200 s | Figure 3 |
| Jump diffusion coefficient | $2.8 \times 10^{11} \text{ nm}^2/\text{s}$ | Section 3.3 |
| Minimum waiting time | Set at 1 ms | Section 2.3 |
| 'Jump' time | Set at 1 ps |  |

The initial pre-synaptic recruitment of Ku70/80 and DNA-PKcs is well reproduced by the combination of mechanisms in our model. The observed behaviour of DNA-PKcs recruitment can be well explained simply as a sequential recruitment of Ku70/80 followed by DNA-PKcs. This makes sense structurally as it has been shown that DNA-PKcs binds to the C-terminal motif of Ku80 (Hammel *et al*, 2010), and Ku70/80 is commonly reported to be the first NHEJ protein recruited to break sites (Ma *et al*, 2004; Lieber, 2010; Chiruvella *et al*, 2013; Wang & Lees-Miller, 2013; Brown *et al*, 2017). The inhibition of Ku70/80 in our model represents competition by alternative pathways for exposed double strand break ends. This means that the rate of Ku70/80 phosphorylation, and subsequent dissociation of the synaptic complex, governs the rate at which double strand breaks have the opportunity to progress down alternative pathways, as suggested by Lee *et al.* (Lee *et al*, 2015).

Due to the self-normalised nature of the experimental data reported, determining the mechanisms governing Ku70/80 recruitment by fitting parameters in our model is challenging and ambiguous. There are conceivably different combinations of the 3 time constants used which would lead to the same overall kinetics. Furthermore, the difference in Ku70/80 recruitment reported by Andrin *et al.* (Andrin *et al*, 2012) and Hartlerode *et al.* (Hartlerode *et al*, 2015) could be explained by either a decrease or substantial increase in the time constant for release from inhibition. Decreasing the time constant would logically result in an increase in the overall recruitment rate of Ku70/80. A substantial increase in the time constant would result in the inhibited state not being released during the time frame investigated and thus the self-normalised Ku70/80 recruitment would represent only the rapid kinetics of Ku70/80 recruitment, unperturbed by inhibition. More rigorous fitting therefore requires a better understanding of the initial mechanisms governing competition by more directly comparable experiments, such as those by Mao *et al.* (Mao *et al*, 2009,2008).

The mechanisms included in this work from Graham *et al.* (G), Lee *et al.* (L), and Uematsu *et al.* (U) combine to produce a constant exchange of DNA-PK at double strand break sites. Specifically these mechanisms combine as follows: Ku70/80 (L) and DNA-PKcs (U) needs to be phosphorylated in order to dissociate. DNA-PKcs, once autophosphorylated in a synaptic complex, is responsible for phosphorylation of Ku70/80 (L). These synaptic complexes are prone to dissociation, but can alternatively stabilise retaining the phosphorylated DNA-PK on the double strand break (G). In this work we have demonstrated that such a combined system is capable of reproducing the observed DNA-PKcs foci kinetics during FRAP experiments as well as the reported long term recovery of ^~^60% pre-bleach intensity (Uematsu *et al*, 2007; Davis *et al*, 2010). Sequential fitting of time constants governing DNA-PKcs recovery was possible due to individual parameters dominating largely separate areas of the FRAP curve, again avoiding over-fitting. Long term recovery was controlled by the rate at which unstable phosphorylated DNA-PK synaptic complexes were stabilised. This could be the result of XRCC4 recruitment by Ku70/80, which together with XLF has been shown to form bridging filaments resistant to dissociation (Reid *et al*, 2015; Brouwer *et al*, 2016). Experiments investigating co-localisation of fluorescently labelled XRCC4 and DNA-PKcs foci would enable implementation and verification of a more detailed mechanistic description of this process in our model.

Having fitted the explicit mechanisms of Ku70/80 and DNA-PKcs recruitment kinetics we show that in order to subsequently fit the general mechanism of repair kinetics for the initial 30 minutes, all fitted time constant are set such that their influence is restricted to the first two hours. In our model therefore, the yield of residual breaks at 24 hours is predominantly determined by the mobility of the individual double strand break ends. This leads to the conclusion that the determining factor which leads to a residual break is the capability of the individual ends to escape the local volume and end up isolated. This mechanism alone is enough to reproduce the LET response of residual breaks at 24 hours observed by Chaudhary *et al.* (Chaudhary *et al*, 2016). However, fitting in this manner causes the model to deviate from experimental data for 1 Gy of 13.7 keV/μm proton irradiation over the time period of 30-300 minutes post irradiation. At 30 minutes and at 360 minutes post irradiation the difference between the Chaudhary *et al.* (Chaudhary *et al*, 2016) data and our simulation is not statistically significant, with p=0.27 and 0.24 respectively via Welch’s t-test. At the intervening points of 60 and 120 minutes, however, the difference is significant with p=0.00001 in both cases. Furthermore, when comparing to a lower LET of 1.77 keV/μm, Chaudhary *et al.* report that repair kinetics show a decreased rate of resolving breaks over that of 13.7 keV/μm radiation during this period, which is the inverse of the simulated behaviour. The behaviour shown in experiment cannot be explained by the differing repair kinetics of complex double strand breaks (Li *et al*, 2014) as high-LET radiation is expected to cause more repair retarding complexities (Henthorn *et al*, 2017). This suggests that there is a missing mechanism behind this behaviour.

Our earlier work showed that the proximity of breaks increases as LET is increased for the same particle type (Henthorn *et al*, 2018). We propose that this LET dependent density could explain the inverse relationship of residual break yield with LET at time points between 30 and 300 minutes. At lower densities, a break that escapes the local area is less likely to encounter another break end and be repaired. This increase in separation from initial partner can be achieved by increasing the mobility of the break end. However, as shown earlier, this would increase the residual yield reported at 24 hours which is currently fitted to experiment. Therefore some mechanism is needed by which breaks have the high mobility necessary to delay formation of synaptic complexes, but remain local enough that given sufficient time meeting another break is probable.

In this work we have chosen to use a CTRW description of sub-diffusion as observed experimentally by Bronstein *et al.* (Bronstein *et al*, 2009) and Girst *et al.* (Girst *et al*, 2013). We show that this leads to sub-diffusive behaviour similar to that reported by others (Bronstein *et al*, 2009; Weber *et al*, 2010; Girst *et al*, 2013; Cabal *et al*, 2006). Having fitted motion to residual double strand breaks from work by Chaudhary *et al.*, it is interesting to note that we also reproduce the ^~^100 nm displacements of individual double strand break ends observed by Soutoglou *et al.* (Soutoglou *et al*, 2007). Soutoglou *et al.* reported that when Ku80 was missing this displacement increased to >500 nm, which could be explained by a dependency of the constrained motion on synapsis formation. If this is the case then it lends weight to the CTRW description of sub-diffusion, which could arise due to the mechanism of transient synaptic complex formation proposed by Graham *et al.* (Graham *et al*, 2016).

In contrast to the CTRW model of sub-diffusive motion, Lucas *et al.* and Weber *et al.* observe behaviours indicating a fractional Langevin motion (fLm) form of sub-diffusion (Lucas *et al*, 2014; Weber *et al*, 2010). This form of motion arises when an object is trapped by a visco-elastic boundary; collisions with which results in the frequent reversals of direction observed by these authors. Lucas *et al.* propose that the surrounding nuclear structures provide this visco-elastic boundary, confining double strand breaks to a local volume. The general diffusion coefficient was found to be 2.4×10^−3^ μm^2^/t^0.5^, two orders of magnitude larger than in this work, and the anomalous diffusion exponent of 0.5, similar to that of this work, whilst the confinement radius was found to reproduce experimental data best at 500 nm. We propose then that this mechanism could explain the inverse relationship of residual break yield with LET at time points between 30 and 300 minutes. The increase in diffusion coefficient would increase the separation of initial partner double strand break ends, whilst the boundary radius would maintain proximity such that the ends are likely to encounter each other again.

## 5 Conclusion

In this work we use an *in silico* model to test the compatibility of several experimentally supported mechanisms hypothesised to operate at distinct stages along the pathway. We determine that the mechanisms proposed by Graham *et al.* (Graham *et al*, 2016), Lee *et al.* (Lee *et al*, 2015), and Uematsu *et al.* (Uematsu *et al*, 2007) can be incorporated into a theory of sequential joining of the major canonical NHEJ proteins (Lieber, 2010; Yang *et al*, 2016; Li *et al*, 2014) to reproduce experimentally observed fluorescent foci kinetics. However, we highlight specific biological mechanisms which are not clearly defined experimentally. Furthermore, we investigate the impact of an assumed CTRW mode of sub-diffusion on the pathway. We show that the motion of individual double strand break ends has a determining role in the repair kinetics of our model for mid (30 min) to late (24 h) time points. Although the motion implemented produces similar displacements as reported by Soutoglou *et al.* (Soutoglou *et al*, 2007), and reproduces the LET dependence of residual breaks reported by Chaudhary *et al.* (Chaudhary *et al*, 2016) we show that CTRW sub-diffusion alone is insufficient to reproduce repair kinetics observed between 60-400 minutes post irradiation. This wide time frame over which double strand break motion can enact substantial influence over the repair kinetic curve highlights an important need for scientific investigation. Improper modelling of this motion could lead to behaviours which need to be compensated for by fitting of the recruitment and activity of repair proteins. This in turn would lead to erroneous conclusions about the kinetics of these proteins and the biological mechanisms they affect.

The model described in this work is incorporated into the existing Geant4 Monte Carlo software familiar to the field of radiation research. As such it can be readily combined with simulations defining the irradiating apparatus and extending these physics models into the biological realm. The model has been constructed in such a way that it can be easily expanded to include relevant biological processes on the scale of individual cell nuclei. Due to following the conventions and structure of Geant4 it can be readily modified and incorporated into existing code by researchers familiar with the tool-kit. As an example we have, in collaboration with Massachusetts General Hospital and the Harvard Medical School, demonstrated the successful implementation of this model into the TOPAS-nBio software (Perl *et al*, 2012; Schuemann, 2012; McNamara *et al*, 2017). These properties make this model a good foundation for continued development by the community towards a multi-scale system capable of clinically relevant simulations.

## Acknowledgements

JWW, NTH, and KJK would like to acknowledge financial support from EPSRC (grant No.: EP/J500094/1). JWW, NTH, KJK, MJM, and RIM would like to acknowledge financial support from the European Commission (grant No.:EC/730983). This research was funded by the STFC Global Challenge Network+ in Advanced Radiotherapy and EPSRC Grand Challenge Network+ in Proton Therapy and supported by the NIHR Manchester Biomedical Research Centre; KJK and MJM would like to acknowledge financial support from the NIHR Manchester Biomedical Research Centre: Radiotherapy Stream (grant No.:BRC-1215-20007). NK and SF would like to acknowledge financial support from the EPSRC (grant No.:EP/N018060/1). MS, MJM, and KJK would like to acknowledge financial support from the European Commission (grant No.:EC/317169). The computational simulation of DNA damage by NTH was achieved using the Condor High Throughput Computing facility at the University of Manchester.

## Author Contributions

JWW developed the DNA repair model, generated, fitted, and analysed the data presented in this work. NTH generated the DNA damage data used in this work. JWW, NK, and SF developed the sub-diffusive motion model. JWW drafted the manuscript based on discussions with MS, RIM, KJK, and MJM. All authors reviewed and approved the manuscript.

